# Stereotaxic targeting of the Dorsal Vagal Complex

**DOI:** 10.64898/2026.06.01.729222

**Authors:** Barbara Robar, Holly E. Smith, Lora K Heisler, Beatrice M Filippi, Pablo B. Martinez de Morentin

**Affiliations:** School of Biomedical Sciences, Faculty of Biological Sciences, University of Leeds, Leeds, UK; The Rowett Institute, School of Medicine, Medical Sciences and Nutrition, University of Aberdeen, Aberdeen UK

## Abstract

The Dorsal Vagal Complex (DVC) is a critical brainstem relay for visceral sensory information, sympathetic regulation, and gut–brain communication. Current weight-reducing pharmacotherapies are reported to target this brainstem region to elicit their main satiety actions. Despite its importance, no published step-by-step protocol exists for stereotaxic targeting of this region in rodents. Here, we present a detailed protocol for bilateral administration of substances into the DVC of mice using the atlanto-occipital membrane approach. We describe the surgical access, obex-referenced coordinate system, injection parameters, and we provide a histological validation. This protocol is useful for the study of DVC cells and efferent and afferent neuronal DVC circuits using common neuroscience tools such as tracings, optogenetics or chemogenetics.

For complete details on the use and execution of this protocol, please refer to Martinez de Morentin et al.(Martinez De Morentin et al., 2024)

## Before you begin

The Dorsal Vagal Complex (DVC) is the primary central relay for vagal afferent information from the gastrointestinal tract, cardiovascular system, and respiratory organs. It is comprised of the nucleus of the solitary tract (NTS), the area postrema (AP) and the dorsal motor nucleus of the vagus (DMX). Subpopulations of the DVC neurons regulate feeding behavior, blood pressure, respiration, and metabolic homeostasis. Recent studies have identified numerous cell-type-specific populations within the DVC through their direct manipulation including, brain-derived neurotrophic factor (Feetham et al., 2024), pre-proglucagon (Jiang et al., 2026), glucagon-like peptide receptor 1(Huang et al., 2024), calcitonin receptor (Cheng, Gonzalez, et al., 2020), cholecystokinin (D’Agostino et al., 2016), catecholamine (Aklan et al., 2020), leptin receptor (Cheng, Ndoka, et al., 2020), pro-opiomelanocortin (Zhan et al., 2013) and astrocytes (MacDonald et al., 2023).

We have recently demonstrated that GABAergic neurons in the DVC (GABA^DVC^) project to the hypothalamic arcuate nucleus (ARC) and regulate food intake and body weight via direct inhibition of NPY/AgRP neurons (Martinez De Morentin et al., 2024). We followed a stereotaxic surgical intervention to reach the DVC that enabled several circuit-based interrogations to define the role of this neuronal subpopulation in feeding behavior and body weight. Specifically, we manipulated Vgat^DVC^ cells using chemogenetic activation and silencing (hM3Dq and hM4Di), visualized Vgat^DVC^ axonal projections using channel rhodopsin assisted circuit mapping (CRACM) and performed specific manipulation of Vgat^DVC-ARC^ axonal terminals using opsins (ChR2).

### Innovation

The stereotaxic delivery of viral vectors into the DVC of mice is technically challenging since it is positioned below the cerebellum (**Figure 1**) and follows a fundamentally different protocol from standard stereotaxic injections. First, the DVC is located on the dorsal surface of the brainstem, requiring access through the posterior fossa rather than through the skull vault. This necessitates flexion of the head and dissection of the atlanto-occipital membrane. Second, the NTS is a small, elongated nucleus flanked by the AP dorsally and the DMX ventrally, creating a narrow corridor for accurate targeting where off-target delivery has significant functional consequences. Third, unlike bregma-referenced coordinates used for brain targets, DVC injections require the obex (*calamus scriptorius*) as the stereotaxic zero-point, a surface landmark that must be visually identified under magnification after membrane dissection. This procedure enables a low off-target rate of the DVC. Although, there are reports using a the atlanto-occipital approach (Joshi et al., 2022), they only provide details to target ventral brainstem regions such as the hypoglossal nucleus or the glomerular reticular nucleus. Given the increasing interest of accessing the DVC to study gut–brain signaling, sympathetic control, and energy homeostasis, we provide a validated step-by-step protocol that allows targeting this brainstem region.

**Figure 1.**
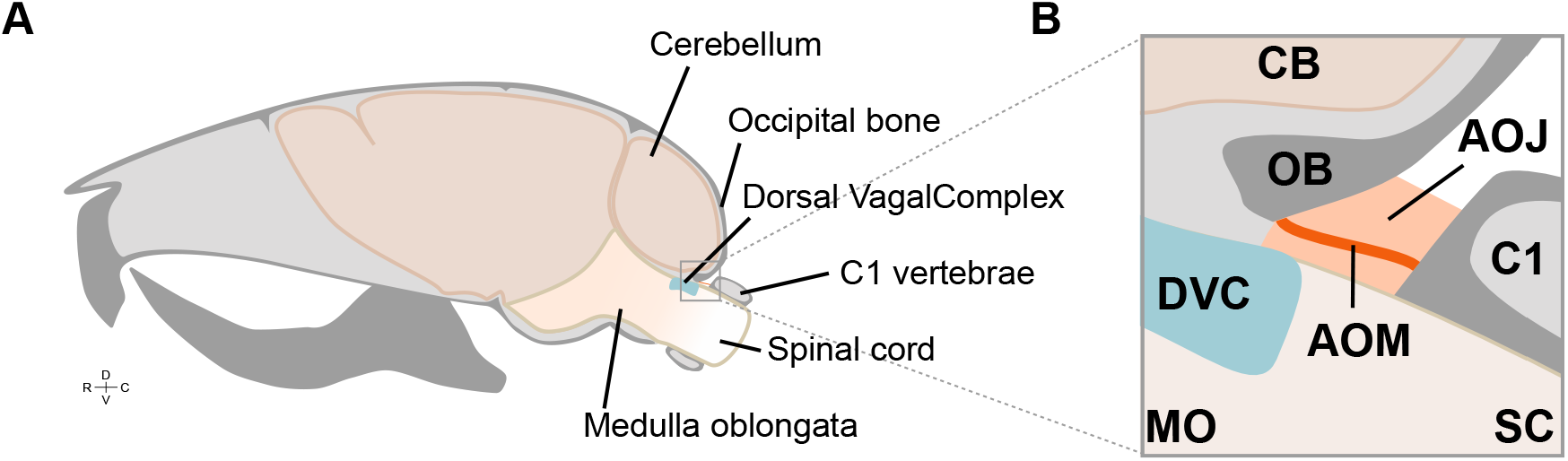
Location of the atlanto-occipital membrane. **A)** Sagittal view of a mouse skull and brain depicting the cerebellum, occipital bone, cervical vertebrae C1, medulla oblongata and dorsal vagal complex. **B)** Representation of the atlanto-occipital membrane spanning the atlanto-occipital joint between the base of the occipital bone and the anterior part of the C1. AOJ: Atlanto-Occipital joint, AOM: Atlanto-Occipital membrane, CB: Cerebellum, C1: Cervical vertebrae, DVC: Dorsal Vagal Complex, MO: medulla oblongata, OB: Occipital bone and SC: spinal cord.

### Institutional permissions

Animal handling and experimentation was carried out according to UK Home Office guidelines and the requirements of the United Kingdom Animals (Scientific Procedures) Act 1986 and the University of Leeds animal welfare ethical review board. Mice were housed under a 12:12 h light/dark cycle with free access to food and water. Two 12-weeks-old *Vgat*^*Cre*^ (JAX016962) male mice were used for this protocol.

***Note:*** *Before applying this protocol in live animals, appropriate ethical approval must be obtained from relevant institutional and government authorities*.

### Preparation of administering agent

#### Timing: 15 min

1. Validation agents. We recommend the use of cresyl violet solution for target validation on post-mortem animals. Cresyl violet allows rapid visualization of targeted region accuracy using a brightfield microscope and enables the visualization of the administered agent spread using 550nm filter in fluorescence microscope.
2. Experimental agents (e.g., viral particles). A viral titration and expression study must be performed to define the optimal conditions.

### Preparation of glass capillary

#### Timing: 20 min

1. Pull glass capillaries using a micropipette puller to achieve a tip diameter of approximately 40μm. (e.g. using Narishige GD1 capillary on a Sutter Instruments P-97 puller: Heat=600, Filament=4, Velocity=60, Delay=145, Pull=175.)
2. Inspect the tip under a microscope to verify diameter and confirm the tip is not broken or jagged.
3. Whether using an automatic or manual injector, we advise marking the capillary with graded lines of volumes.

***Note***: *If using G1 capillary (0D=1mm, ID=0*.*6mm), a 1mm mark interval correspond to approximately 250nl*.

4 Fill the capillary with the administering agent by gentle aspiration using an microinjector (e.g. pneumatic IM-11-2 Narishige).
5 Test-eject a small volume to confirm flow before mounting on the stereotaxic arm.

### Preparation for surgery

#### Timing: 30 min

This section is detailed as if the procedure would be performed in live animals under general anesthesia, but training should be performed on animals post-mortem.

1. Surgery sets
  a. A set of standard surgical tools (Figure 2) including:
    i. Micro-dissecting scissors
    ii. Forceps
    iii. Mosquito
    iv. Absorption paper
    v. Coton swab sticks
  b. A set of custom dissecting needles (**Figure 3**) that includes:
    i. scalpel
    ii. Hook
    iii. Muscle retractor
    iv. Cerebellum retractor
  c. Surgical gowns and drapes.

**Figure 2.**
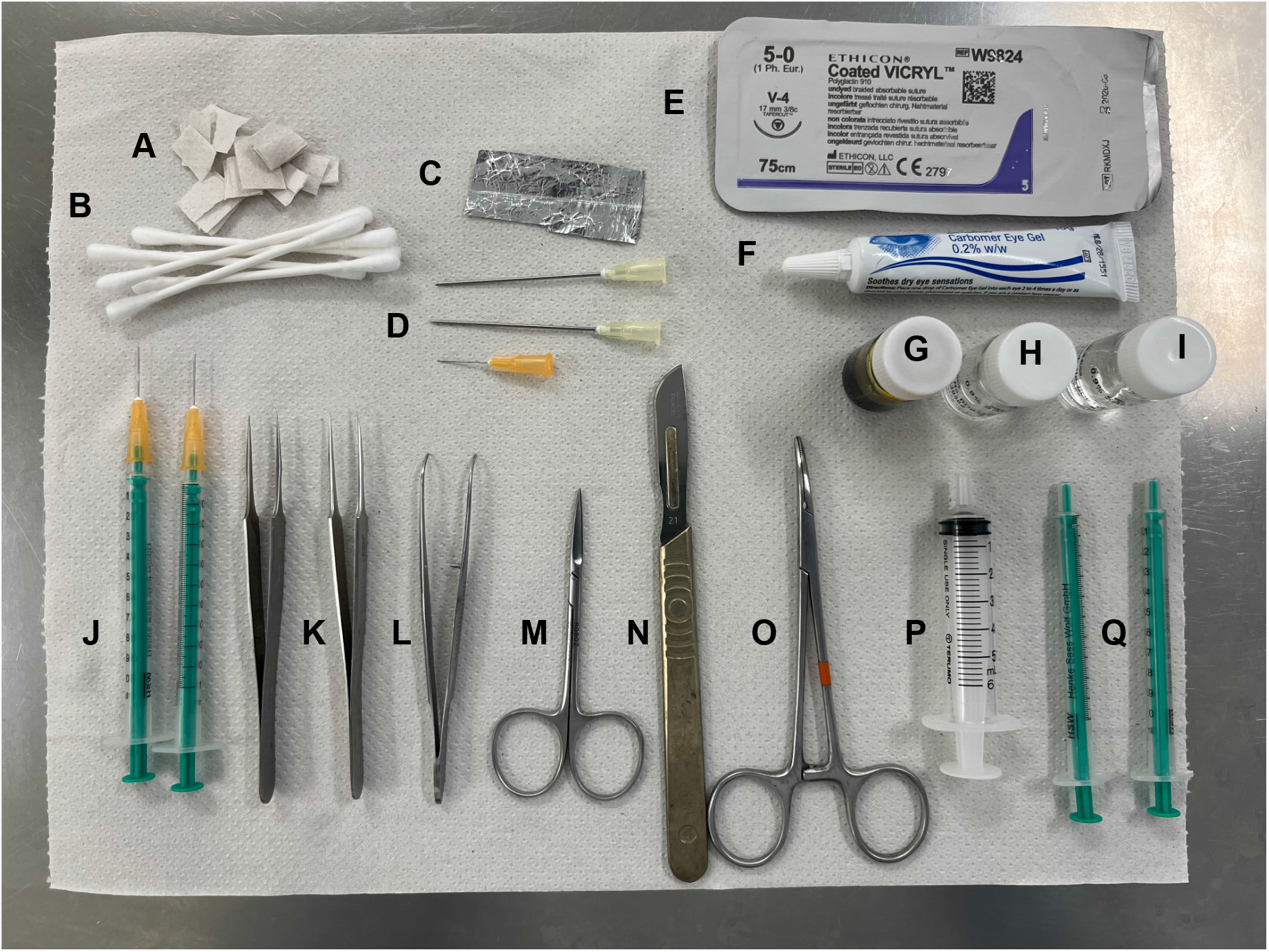
Surgical tools. **A)** Absorbent paper, **B)** Cotton buds, **C)** Graded capillary, **D)** retractor-needles, **E)** Absorbent suture, **F)** Eye lubricant, **G)** Iodine Povidone, **H)** Saline, **I)** Analgesic, **J)** Dissecting needles (scalpel and hook), **K)** Fine dissecting forceps, **L)** Fine dissecting forceps with teeth, **M)** Fine dissecting scissors, **N)** Scalpel, **O)** Fine needle holder, **P)** 5ml syringe, **Q)** 1ml syringes.

**Figure 3.**
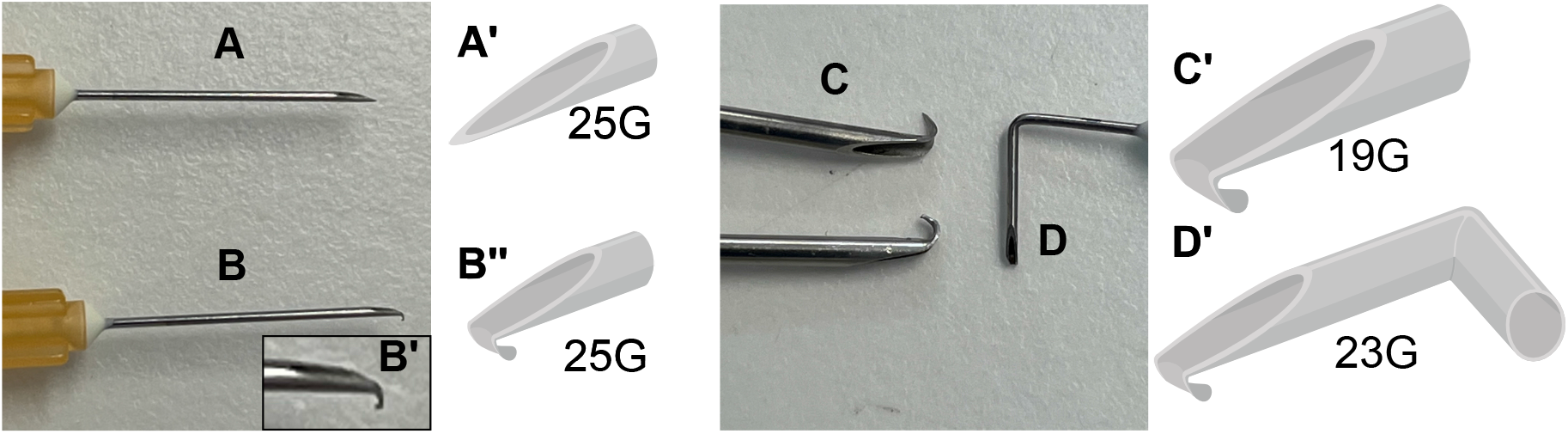
Custom dissecting needles and diagrams. **A)** scalpel, **B)** hook, **C)** muscle retractors and **D)** cerebellum retractor.

***Note:*** Use transparent self-seal sterilization pouches for each reusable set.

2. Animal preparation.
  a. Include animal details in the surgery record’s book.
  b. Set the recovery heating cabinet at approximately 35°C and verify with an independent thermometer.
  c. Prepare new home cages with standard enrichment. Mash standard pelleted diet in warm water (35°C) and place in a petri dish inside the new home cage.
3. Surgical station.
  a. Check all surgical tools are autoclaved, dried, and cool.
  b. Given the unusual positioning of the mouse head (60° forward) rodent anesthetic masks and tooth restrainers on most stereotaxic frames are not suitable for DVC surgery. We have 3D printed a mask that accommodate the variable angle of the head (**Figure 4A**). Similar anesthetic masks can be obtained from specialist suppliers.
  c. Confirm adequate anesthetic and oxygen supply and retrieval thought the mask.
  d. Pre-fill a 1 mL syringe with warm (35°C) sterile 0.9% saline; place on a heat mat (35°C) (for surgical subcutaneous rehydration).
  e. Pre-fill a 5 mL syringe with sterile saline; place in a beaker on wet ice for bleeding control and wound cleaning.
  f. Clean the stereotaxic frame and surgical area with 70% ethanol.
  g. Place sterile surgical drapes.
4. Surgeon gowning and preparation.

**Figure 4.**
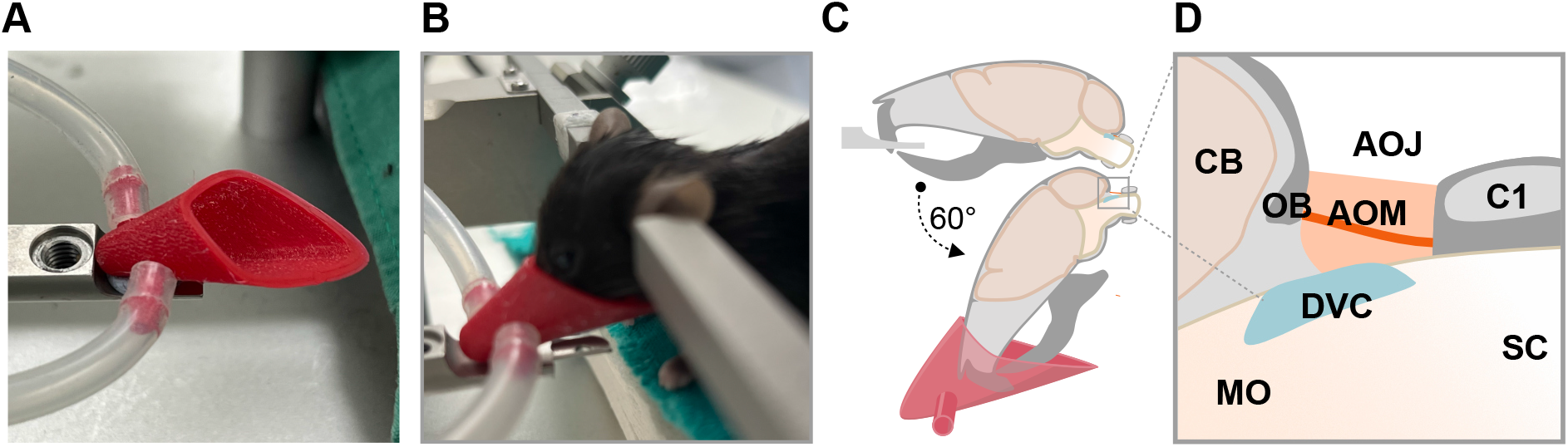
Angled head with custom anesthetic mask. **A)** 3D printed anesthetic mask. **B)** representative position of the mouse head angled. **C)** Diagram of the head angled 60°. **D)** Diagram of the atlanto-occipital joint with the head angled. AOJ: Atlanto-Occipital joint, AOM: Atlanto-Occipital membrane, CB: Cerebellum, C1: cervical vertebrae 1, DVC: Dorsal Vagal Complex, MO: medulla oblongata, SC: spinal cord.

Follow standard aseptic gowning procedure and principles (e.g. LASA, RAT), as described elsewhere (Imoesi et al., 2026).

5. Induction of anesthesia.
  a. Place the mouse in the pre-anesthetic chamber and set the anesthetic flow (3-5% isoflurane in oxygen).
  b. Once fully anesthetized as illustrated by the lack of pedal withdrawal reflex, transfer the mouse to the stereotaxic frame and place the nose in anesthetic the mask.
6. Maintenance of anesthesia and pre-operative preparation.
  a. Maintain isoflurane at 1–2% in oxygen throughout the procedure.
  b. Secure head to the stereotaxic frame using ear bars.
  c. Clean fur and disinfect with cotton wool with povidone-iodine. **Note:** we advise a thorough cleaning and disinfection without the need of shaving to reduce the risk of surgical site infection due to micro-abrasions and skin cuts.
  d. Apply eye hydration (e.g. carbomer, directly to the eyeball) and provide light protection.
  e. Apply analgesic and anti-inflammatories.
  f. Apply pre-warmed (35°C) hydration fluid saline (1mL/30 g mouse, SC)
7. Positioning of the head

At this step, the DVC protocol diverges from a standard stereotaxic brain surgery protocol. From a position where the skull is horizontal, incline the head forward approximately 60 degrees (**Figure4B-D**). Adjust the mask to the new position and make sure it is sealed on the mouse’s snout.

**CRITICAL:** The head flexion angle is critical. If the angle is low, the atlanto-occipital membrane will not be accessible. If too high, the mouse’s airway may become obstructed.

**Tip:** Once the head is flexed, if a click is heard with each of the mouse’s inhalation, reduce the angle immediately. This noise signifies that the trachea is being squeezed by the head angle. It is essential to monitor breathing continuously.

### Step-by-step procedure details

#### Exposure of the dorsal brainstem

##### Timing: 10-15 min

This section describes the surgical approach to expose the dorsal brainstem surface for DVC access. This approach is fundamentally different from a standard stereotaxic protocol and requires practice with cadaver tissue before undertaking on live animals. The entire process is performed using a stereo microscope.

1. Exposing the posterior atlanto-occipital membrane. *Here we aim to dissect the dermis to expose the atlanto-occipital joint*.
  a. Looking at the back of the mouse’s neck, identify a flat horizontal area in the skin (**Figure 5A**). This flat region is due to the distension of the neck stretching the skin between the skull bone and the C1 vertebrae.
  b. Pinch the skin with the forceps, lift the skin and cut it longitudinally with the pair of surgical scissors to create a 0.5 to 1cm incision. This will reveal the hypodermis (**Figure 5B and Supplementary Video 1**). **Tip:** We will use retractors, so as little as 0.5 cm would be sufficient for a skilled surgeon and will accelerate the wound recovery.
  c. Cut the hypodermis following a process as detailed in section b. This will reveal the right and left neck muscles (semispinalis capitis) running from the base of the occipital bone to the cervical vertebrae C1. A white pale midline is visible between the muscles (**Figure 5C and Supplementary Video 2**).
  d. Using the *scalpel* and the *hook needles* cut along the midline and gently pull the left and right muscle apart (**Figure C’ to D’**).
  e. Using the *scalpel needle*, detach the ventral layer of the neck muscles from the membrane and pull the muscle further apart using the retractor. Repeat this process for the other neck muscle (**Figure D’ to E’**).
  f. Secure the separated muscles using *muscle retractors*. This will allow you to visualize the posterior atlanto-occipital membrane (AOM), which is a translucent layer spanning the base of the occipital bone and the C1, protecting the atlanto-occipital joint (AOJ) (**Figure 5E and Supplementary Video 3**).

**Figure 5.**
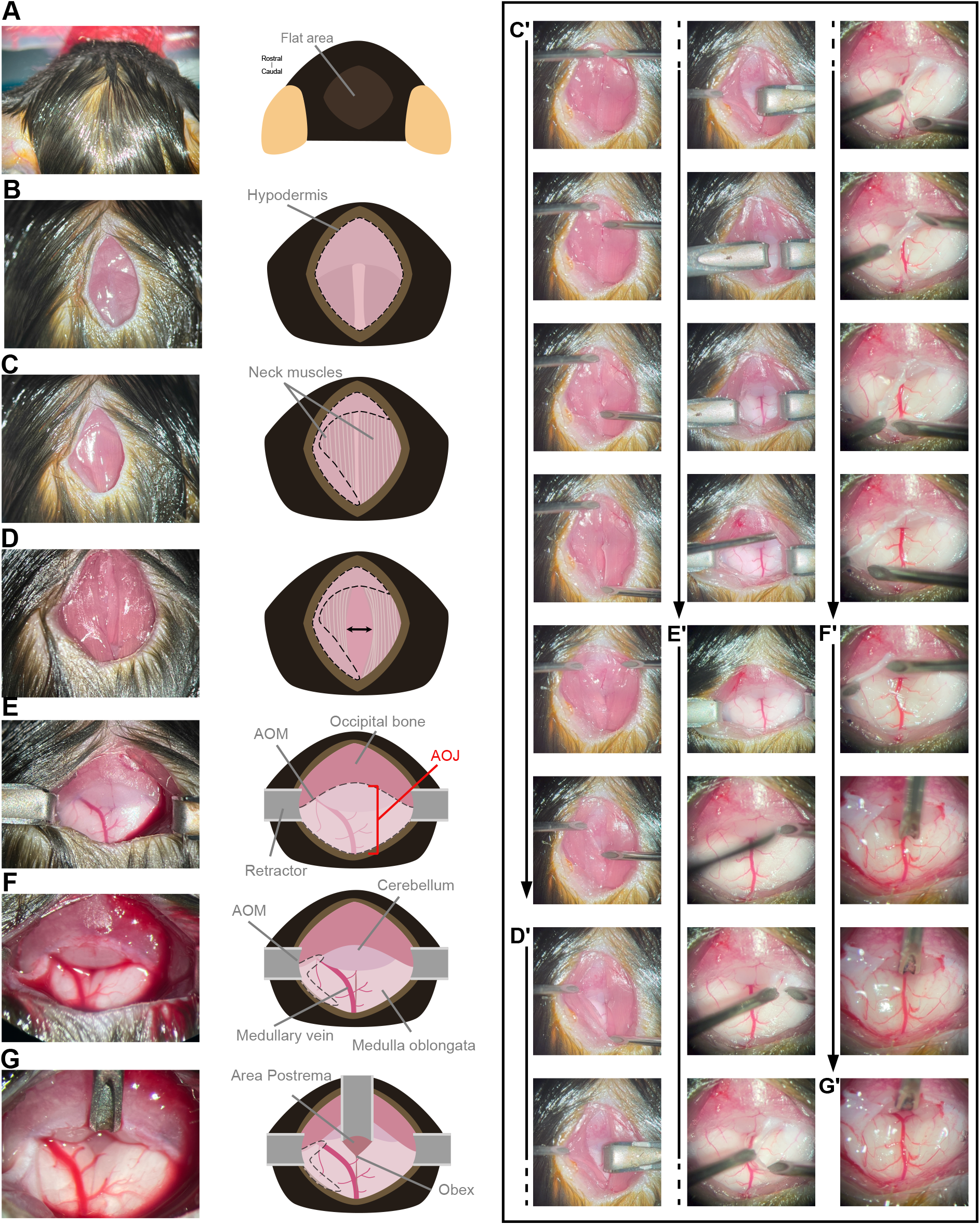
Dissection of the atlanto-occipital membrane and access to the dorsal branstem. **A)** Identification of virtual flat at the back of the neck. **B)** Incision on the skin and exposure of hypodermis. **C)** Dissection of hypodermis and exposure of neck muscles. **D)** Separation of the muscles. **E)** identification of AOM and AOJ. **F)** Dissection of AOM. **G)** Retraction of cerebellum. **C’ to G’)** Panel with detailed dissection images taken from video captures in Supplementary Videos 1 to 7.

**Figure 6.**
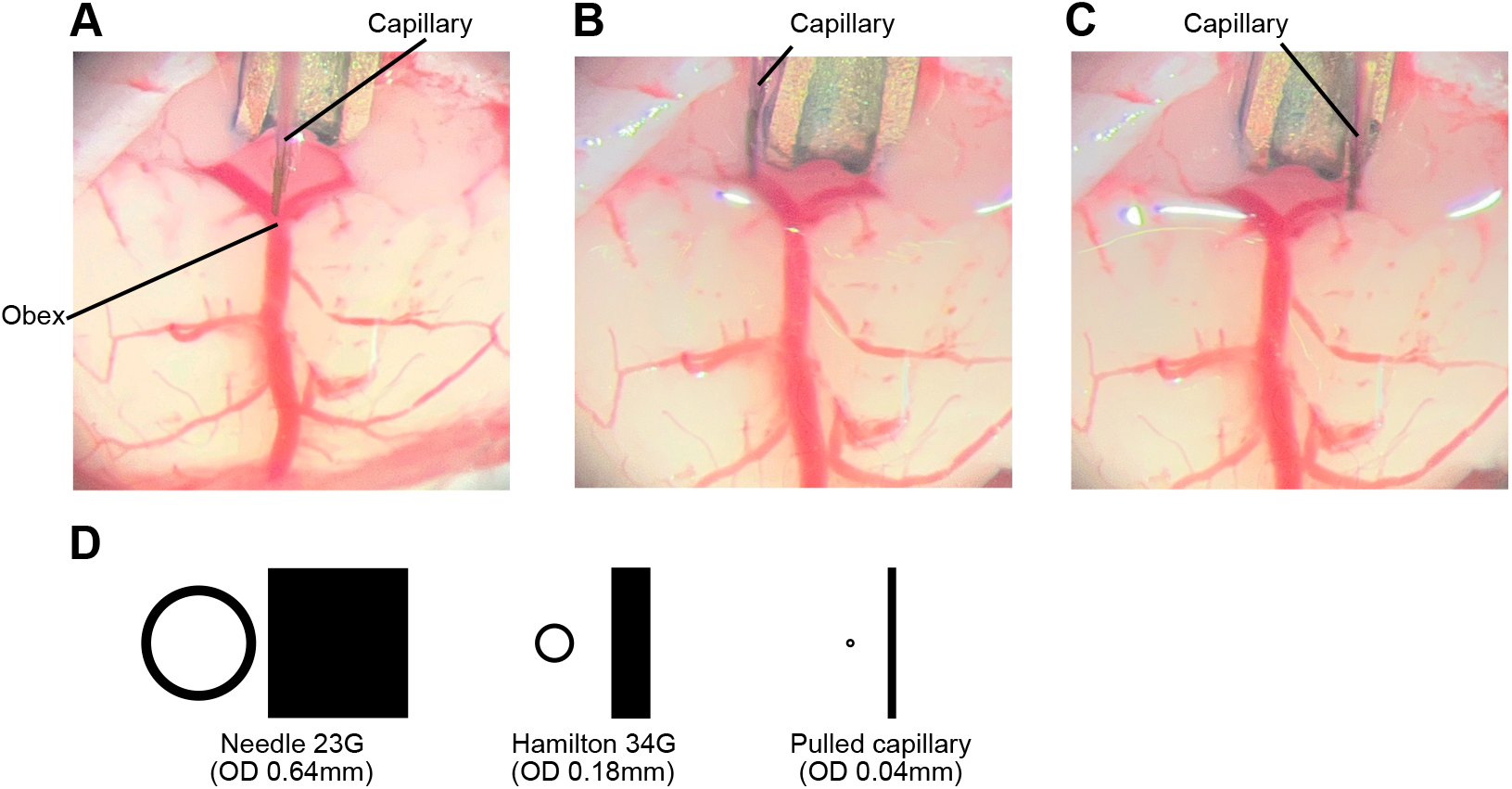
Bilateral administration of injectable agent into the NTS. **A)** Zero of stereotaxic corrdinates placing the tip of the capillary above the obex; **B)** injection on the left side (L:-0.25mm, DV:-0.25mm) and **C)** injection on the rigth side (L:+0.25mm, DV:-0.25mm). **D)** Comparison of diameter and cross section between 23G cerebellum retractor needle, Hamilton 34G and capillary

**CRITICAL:** Given the limited anatomical space, we advise using needles (19G) with their tips bent down as retractors. Their tips must be also blunted to prevent damaging the muscle or tearing the AOM.

***Note:*** *At the end of this stage of the procedure, you should have a 1 cm plane to identify:*

i. Top: Occipital bone (pink)
ii. Middle: ventral cerebellum (pale pink)
iii. Bottom: medulla oblongata (white)
iv. Median posterior medullary vein (red)

2. Dissecting the atlanto-occipital membrane. *Here we want to remove the AOM to access the AOJ*. ***Note:*** *The median posterior medullary vein accesses the skull cavity laterally. The dissection must take place on the opposite side of the vein to prevent any damage to the vein*.
  a. Using the *scalpel needle*, perform a vertical perforation of the AOM at the joint with the occipital bone. This will relief the inner pressure of the AOJ and release CSF, using the *hook needle* and a sharp needle as scalpel, proceed to dissect the AOM caudally and laterally towards the opposite side (**Figure 5E’ to F’ and Supplementary Video 4**). **CRITICAL:** When dissecting close to the vein pull the AOM up using the retractor needle to allow sufficient space for the scalpel to cut the AOM over the vein.
  b. If necessary, gently clean CSF or blood from the area using sterile cotton buds. ***Note:*** *A small degree of bleeding is acceptable as a tiny vessel can be damaged during this part of the procedure. However, if the medullary vein is lesioned, as indicated by bleeding that does not stop, then the mouse may suffer a brain stroke during recovery, we advise to immediately humanely kill the mouse*.
  c. Once the AOM is completely dissected and the inner pressure is relieved, the medulla and the cerebellum will slightly protrude (**Figure 5F**).
  d. Using the *cerebellum retractor needle*, pull the base of the cerebellum rostrally until the obex (calamus scriptorius) is visible (**Figure 5F’ to G’ and Supplementary Video 5**).
  e. The obex appears as a small V-shaped notch at the caudal end of the visible brainstem surface. The area postrema is visible as a slightly raised, vascularized region immediately rostral to the obex. The aim is to reveal an area of approximately 2mm between the obex and the retractor (**Figure 5G**).
  f. Secure the retractor at the desired position. ***Note***: *Adequate illumination and magnification are essential. If the surface is obscured by blood or CSF, gently irrigate with saline and absorb with absorbent paper*.

### Bilateral administration of injectable agents into the NTS

This section describes how to administer an injectable agent into the dorsal brainstem targeting the NTS. The AP and the DMX may also be targeted using this technique.

#### Timing: 10–20 min

1. Affix the graded glass capillary (or a 34G Hamilton needle) to the stereotaxic syringe adapter.
2. Load the injectable agent.
3. Load a small volume of air sufficient to prevent the withdrawal of the agent by capillarity when the capillary enters in contact with the CSF.
4. Position the tip of the capillary above but touching the obex (**Figure 6A**).
5. Zero the stereotaxic coordinates to this position.
6. For targeting the NTS: Lift the capillary slightly and move it to the coordinates: antero-posterior -0.25mm and lateral +0.25mm, to the obex. This should leave the tip of the capillary above the commissure between the AP and the NTS (**Figure 6B**).
7. Displace the capillary ventrally -0.25 mm.
8. Start the administration of 250nl of agent at rate of 50nl/min.
9. Leave the capillary on the target position 5 minutes.
10. Slowly retract the capillary.
11. Move the capillary laterally to the opposite side -0.25 and repeat the injection (**Figure 6C and Supplementary Video 6**).

***Note:*** *Microliter syringes can be used, but we recommend using glass capillary pipettes. We provide a cross-section comparison of the thinnest available needle (Hamilton, 34G) and a pulled capillary (GD-1) (***Figure 6D**).

***Note***: *In some mice, the medullary vein will travel into the brain following the commissure of either side. The vein can be gently pushed to a side with the tip of the capillary*.

### Closure of the wound

This section describes how to reposition the semispinalis capitis muscles and suture the skin.

#### Timing: 10 min procedure + post-surgical monitoring time (as needed)

1. Clean the AOJ by gently flushing with sterile saline and remove any clotted blood.
2. Slowly reposition the head of the mouse to an almost horizontal plane.
3. Hydrate the neck muscles by flushing saline and perform a gentle massage with aseptic techniques to return them to position. ***Note:*** *A small drop of surgical glue can aid in re-bonding both muscles together*.
4. Pull the skin together and clean it from inside to outside with a cotton bud with sterile saline.
5. Add stitches of absorbable suture to close the wound. **CRITICAL:** Brainstem surgery carries risks of respiratory depression and swallowing dysfunction. If weight loss exceeds 15% or respiratory distress is observed, apply humane endpoint criteria immediately.

## Expected outcomes

Before performing experimental surgeries, extensively validate coordinates and technique by injecting cresyl violet dye into a mouse cadaver. Successful targeting is confirmed when dye is localized to the medial NTS with minimal spread into the AP or DMX.

For experimental animals, successful viral transduction is indicated by bilateral reporter expression restricted to the NTS, spanning 4–6 coronal sections (∼200– 400μm). We have transduced a *Vgat*^*Cre*^ mouse with viral particles expressing red (AAV-hSyn-DIO-mCherry) and green (AAV-hSyn-DIO-GFP) reporters into the NTS as an example (**Figure 7**).

**Figure 7.**
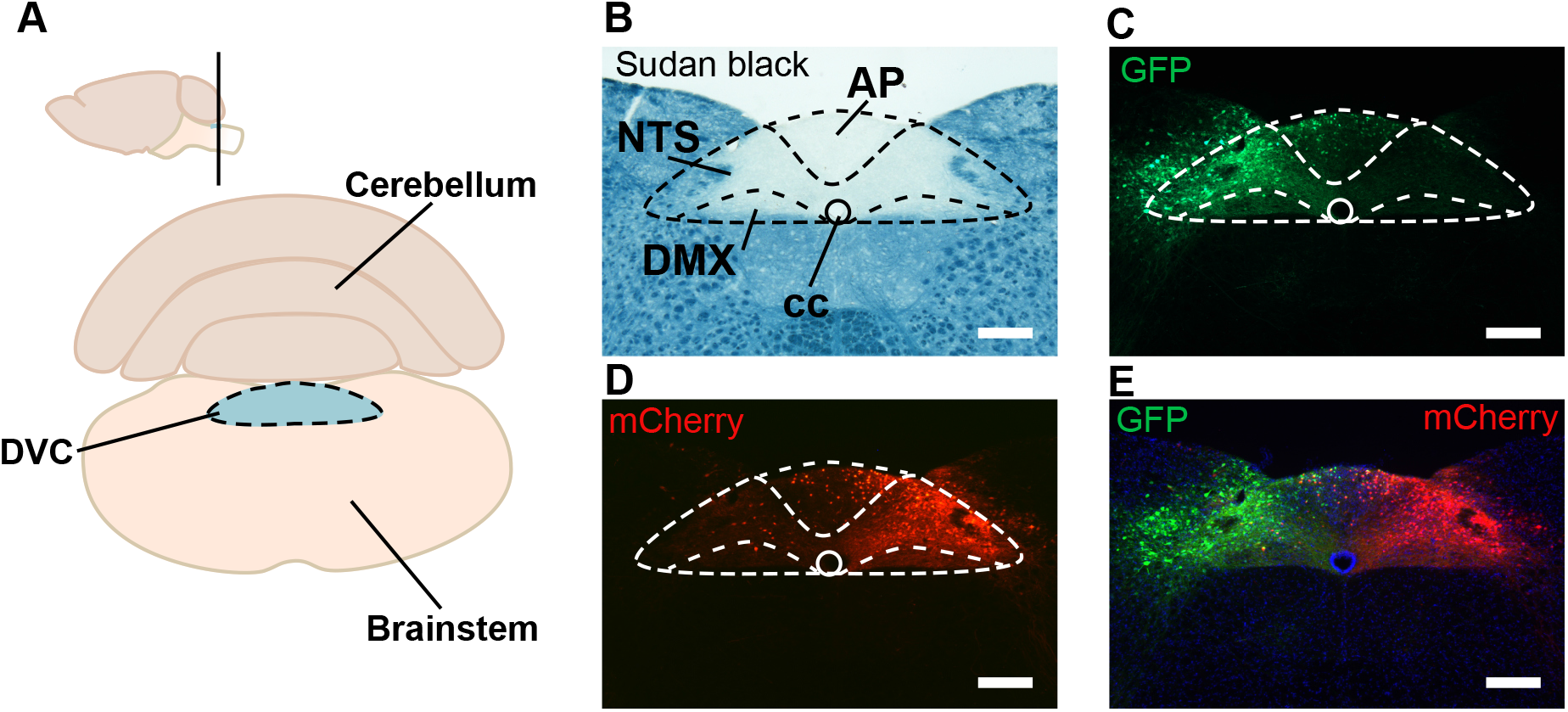
Representative DVC targeting with AAV expressing reporters. **A)** Diagram of coronal section of the brainstem containing the DVC. Photomicrographs of the DVC of a *Vgat*^*Cre*^ mouse **B)** counterstained with Sudan black B, **C)** transduced with AAV-hSyn-DIO-GFP on the left side, D) transduced with AAV-hSyn-DIO-mCherry on the right side and **E)** image merged. Scale bar 200 μm. AP: area postrema; cc: central canal; NTS: Nucleus of the Solitary Tract; DMX: Dorsal Motor Nucleus of the Vagus.

The accuracy of DVC injection has been further validated by functional readouts published (Martinez De Morentin et al., 2024).

## Limitations

We acknowledge that there exist several limitations of technical nature when performing this surgical procedure.

The primary limitation is the surgeon’s skills. A critical step comprises the dissection of the AOM without damaging the medullary vein. This can otherwise lead to the need to humanely kill the animal. It is essential to practice technique competency in cadavers before moving to live animals.

The DVC is an elongated structure, and it is formed by three main structures: the AP, the NTS and the DMX. If your study design uses an intersectional approach (e.g. Cre/LoxP technology) we recommend carefully reviewing the off-target possibilities. We highly recommend optimizing: 1) agent titer/concentration, 2) the volume of the agent administered and 3) adapting the coordinates from the obex.

The transduction volume is constrained to small amounts. Exceeding 300nl risks off-target transduction. For broader coverage, multiple injection sites along the AP axis may be considered.

## Troubleshooting

### Problem 1

Mouse with dyspnea and/or gasping (clicking noise)

### Potential solution

Reduce the concentration of anesthetic. Lower the angle of the head. Prevention: verify consciousness and angle the head slowly.

### Problem 2

Excessive bleeding during membrane dissection

### Potential solution

Apply cold sterile saline; wait 1–3 min. Apply gentle pressure with absorption triangles for 3-5 min. If from the medial posterior medullary vein, consult the animal welfare officer. Prevention: use only blunt dissection needles, avoid lateral excursions, ensure adequate visualization.

### Problem 3

Inability to identify the obex

### Potential solution

Irrigate with room-temperature sterile saline and absorb with sterile cotton buds. Improve illumination and magnification. Verify head angle (∼60°). Use a deeper retractor for the cerebellum. Lower the thorax. If the obex cannot be identified, do not proceed and humanely kill the animal.

### Problem 4

Blocked capillary

### Potential solution

Given how superficial the target region is, the tip of the capillary can be cut several times if it is blocked while maintaining its diameter.

### Problem 5

Medullary vein above the commissure AP-NTS

### Potential solution

Use a blunt fine tool to displace it during the positioning of the capillary. Use the capillary to push the vein to the side.

### Problem 6

Off-target expression of the agent in the brain

### Potential solution

Reduce volume of agent. Reduce the administration rate. Increase micropipette insertion time to 10 min. Re-adjust the coordinates.

### Problem 7

Post-operative respiratory distress or mortality

### Potential solution

This may indicate brainstem trauma. Ensure DV injection is conservative (−0.25 mm max). Do not exceed 300nl of agent infusion. Maintain mouse body normal temperature and hydration.

### Problem 8

Wound re-opening

### Potential solution

When suturing, place the first knot close to skin with two additional knots. Add a drop of surgical glue to the knot. Re-suture once if stitches are removed within the first 24 h. If beyond 24 h or infected, apply humane endpoints. This may be more recurrent in female mice due to social grooming behavior. One option is to single house female mice for 48h until wound has healed.

## Resource availability

### Lead contact

Further information and requests should be directed to Pablo B. Martinez de Morentin.

### Technical contact

Technical questions should be directed to Pablo B. Martinez de Morentin.

### Materials availability

This protocol uses generated 3D printed models that are available upon request.

### Data and code availability

Representative histological data are presented in this protocol. Additional data available from the lead contact upon request.

## Acknowledgements

This work has been supported by University of Leeds PhD studentship to B.R.; Yorkshire Bioscience DTP to H.E.S.; UKRI-BBSRC (BB/V016849/1) to L.K.H.; MRC-Career Development Fellowship (MR/S007288/1) and Wellcome Trust (UNS63234) to B.M.F. and University of Leeds (start-up package) and The Royal Society (RGS\R2\252055) to P.B.M. We thank Dr. Cristian Olarte-Sanchez and Dr. Guisseppe D’Agostino for their advice and staff at Centre of Biomedical Services, University of Leeds for their assistance.

## Author contributions

Conceptualization, P.B.M. and B.M.F.; methodology, P.B.M and B.M.F.; writing - original draft, P.B.M; writing - review and editing, B.R. H.S.; L.K.H.; B.M.F. and P.B.M.; funding acquisition, P.B.M. and B.M.F.; resources, P.B.M. and B.M.F.; supervision, and project administration, P.B.M.

## KEY RESOURCES TABLE

| REAGENT or RESOURCE | SOURCE | IDENTIFIER |
| --- | --- | --- |
| **Chemicals, peptides, and recombinant proteins** |  |  |
| Isoflurane | MERCK | 792632 |
| Carprofen | Rimadyl™, Zoetis |  |
| Sterile 0.9% sodium saline | Cambridge Bioscience | TSH-00526 |
| Iodine-povidone solution | Vetasept® Povidine-Iodine Antiseptic Solution. Animalcare | XHG012 |
| Carbomer eye gel 0.2% | Medicom | -- |
| Cresyl violet dye | MERCK | C5042 |
| AAV9-hSyn-DIO-EGFP | Addgene | 50457 |
| pAAV9-hSyn-DIO-mCherry | Addgene | 50459 |
| Ethanol | Fisher Scientific | Cas# 64-17-5 |
| Virucidal disinfectant | Virkon™ | 12338667 |
| **Experimental models: Organisms/strains** |  |  |
| Male mouse: VgatCre | Centre of Biomedical Services | IMSR_JAX:016962 |
| **Other** |  |  |
| Micro-dissecting scissor, size 4 | MERCK | S3146 |
| Micro-dissecting forceps | MERCK | F4017 |
| Needle Holder | WPI | 14110 |
| Absorbent paper | Whatman | 1440329 |
| Cotton buds | Camlab Limited | 1234602 |
| Stereotaxic Instrument (Kopf 942) | Kopf | 900SL |
| Pneumatic Microinjector IM-11-2 | Narishige | IM-11-2 |
| Glass Capillaries | Narishige | GD-1 |
| Pipette puller | Sutter Instruments | P-97 |
| Micro syringe | Hamilton Bonaduz | 7632-01 |
| Removable needles 34G | Hamilton Bonaduz | 207434 |
| Suture 6/0 absorbable | Ethicon | ETHW9913 |
| Needle 19G | Greiner Bio-One Ltd | N1950 |
| Needle 25G | Greiner Bio-One Ltd | N2516 |
| Needle 23G | Greiner Bio-One Ltd | N2316 |
| Syringe 1ml | SLS | SYR6200 |
| Syringe 5ml | SLS | SYR6204 |
| Stereo microscope | Zeiss |  |

